# Genetic mapping, marker development, and identification of candidate genes for powdery mildew resistance in *Malus baccata* ‘Jackii’

**DOI:** 10.1101/2025.08.26.672377

**Authors:** Matthias Pfeifer, Leonard Kurzweg, Buist Muçaj, Tom Burkhardt, Andreas Peil, Henryk Flachowsky, Ofere Francis Emeriewen, Thomas Wöhner

## Abstract

Powdery mildew, caused by *Podosphaera leucotricha*, is one of the most important fungal diseases in apple cultivation worldwide. *Malus baccata* ‘Jackii’, however, exhibits resistance to this pathogen, and a previous study demonstrated that this resistance co-segregates with an AFLP marker on linkage group 10. The objectives of this study were to construct genetic linkage maps, develop molecular markers, and identify candidate genes associated with this powdery mildew resistance, making use of the *Malus baccata* ‘Jackii’ genome sequence. An F1 population derived from the cross ‘Idared’ × *Malus baccata* ‘Jackii’ was phenotyped from 2009-11 and 2023-24 and genotyped with SNPs generated from tGBS and SSRs. Genetic linkage maps of *Malus baccata* ‘Jackii’ were constructed, and QTL mapping confirmed the presence of the powdery mildew resistance locus *Plbj* on linkage group 10. In addition, a minor QTL was detected on linkage group five. Closely linked SSR and KASP markers were developed, and the *Plbj* locus was delimited to a 3,287,286-bp region on haplotype one of the *Malus baccata* ‘Jackii’ genome, in which resistance gene candidates were identified. This study supports the direct application of molecular markers in breeding programmes and provides an essential background for future functional studies of *Plbj*.

## Introduction

The domesticated apple (*Malus domestica* Borkh.) is an important temperate fruit crop that faces multiple biotic stresses, and changes in climate and pathogen populations may further increase disease impact (Hanke et al. 2020; Strickland et al. 2021). Among the most significant pathogens is powdery mildew, caused by species of the genus *Podosphaera*. These fungi are highly adaptable and can infect over 10,000 plant species with frequent reports of host jumps and host range expansions (Kusch et al. 2024). Consequently, a perpetual race emerges between the fungus and plant breeders, who must persistently monitor and seek novel resistances. *Podosphaera leucotricha* (Ellis & Everh.), the primary powdery mildew pathogen of apple, has also been observed on other hosts such as *Photinia × fraserii*, *Prunus africana* and *Pyrus calleryana* (Garibaldi et al. 2005; Minnis et al. 2010; Mwanza et al. 2001). This obligate biotrophic, ascomycetous, heterothallic, ectoparasitic fungus causes considerable economic losses and increases production costs in apple cultivation each year (Coyier 1974; Takamatsu 2013; Yoder 2000). The mycelium overwinters inside buds and, after bud burst, initiates primary infections, which cause delayed growth and deformations of shoots, leaves, flowers and fruits (Strickland et al. 2021). Sexual reproduction via ascospores is possible but appears to play a minor role in disease spread, whereas the formation of asexual conidia can drive multiple secondary infection cycles within a single season (Strickland et al. 2021). Management strategies to prevent yield losses include crop cultural practices, as well as chemical and biological measures (Strickland et al. 2021). The fungicides currently used in apple cultivation remain effective against the pathogen (Strickland et al. 2023). Their use in consistent rotation is essential to prevent resistance development (Strickland et al. 2023), especially since the emergence of fungicide resistance in powdery mildew has been frequently observed (Lesemann et al. 2006; Vielba-Fernández et al. 2020). The most desirable and environmentally sustainable strategy to control apple powdery mildew is the growing of resistant cultivars. This approach can also make apple production more economical by reducing both the risk of yield losses and the costs of control measures.

Various major resistances to powdery mildew, defined as single genes or clusters of genes with large effects providing qualitative resistance, are present in the genus *Malus*. *Pl-1* and *Pl-2* were identified in *Malus* × *robusta* and *Malus* × *zumi* (Knight and Alston 1968), *P-lw* in ‘White Angel’ (Gallott et al. 1985), *Pl-d* in accession D12 (Visser and Verhaegh 1980), *Pl-m* in Mildew Immune Selection (Bus et al. 2010; Dayton 1977) and *Plbj* in *Malus baccata* ‘Jackii’ (Dunemann and Schuster 2009). Furthermore, powdery mildew resistance has also been reported in the apple clone U 211 (Stankiewicz-Kosyl et al. 2005) and in *Malus florentina* and *Malus sieboldii* (Schuster 2000). Moreover, the knock-down of the expression of *Mildew Locus 0* (*MLO*) genes could have the potential to make apple trees more resistant to powdery mildew, though further research is required to fully understand the mechanisms involved (Pessina et al. 2016; Pessina et al. 2017). As observed in apple, powdery mildew resistance genes can be overcome, as shown for *Pl-1* (Kellerhals et al. 2013; Krieghoff 1995) and *Pl-2* (Caffier and Laurens 2005; Caffier and Parisi 2007). Therefore, the combination (pyramiding) of different powdery mildew resistance genes in a cultivar should be the objective, as this has the potential to increase the durability of the resistance (Mundt 2018). Consequently, it remains of great importance not only to identify additional resistance genes, but also to facilitate the utilisation of existing ones. A previous study reported the presence of the *Plbj* resistance locus on linkage group (LG) 10 in *Malus baccata* ‘Jackii’ (Dunemann and Schuster 2009). The primary objectives of this study were to generate a higher-resolution map of LG 10, further narrow down the genetic region containing the *Plbj* resistance locus, and verify whether *Plbj* is still effective in our experimental field. Therefore, offspring from a cross between ‘Idared’ and *M. baccata* ‘Jackii’ (hereafter referred to as *Mb*j) were genotyped using various marker systems and phenotyped for susceptibility to powdery mildew over several years in the experimental field at the Julius Kühn-Institut (JKI) in Dresden-Pillnitz, Germany. An additional aim was to develop molecular markers tightly linked to *Plbj* for application in marker-assisted selection (MAS) and to identify resistance gene candidates using the haplotype-resolved genome sequence of *Mb*j (Pfeifer et al. 2025, preprint).

## Results

### Phenotyping results of the ‘Idared’ × *Mb*j F_1_ populations

An overview of the phenotyping results is presented in Table 1. In the primary mapping population in the field (122 F_1_ individuals of ‘Idared’ × *Mb*j, designated 05225 and 06228) the highest mean powdery mildew score was observed in summer 2023 (2.76), whereas the lowest mean score occurred in spring 2010 (1.34). Maximum disease severity among the offspring ranged from seven to nine, indicating consistently high infection pressure in the field. Across all years, 71 individuals were classified as resistant, while 51 were scored as susceptible at least once. In 2023 and 2024, only 120 individuals could be evaluated due to the loss of two genotypes. Spearman correlation coefficients between field assessments ranged from 0.61 (summer 2010 vs. spring 2011) to 0.97 (spring 2023 vs. spring 2024). Phenotypic distributions consistently showed a right-skewed pattern. In the secondary F_1_ population (127 individuals, designated 24230) from the same cross combination, phenotyped in the greenhouse in 2025, 46 individuals were resistant and 81 were susceptible. Considering both populations, a total of 117 were classified as resistant and 132 as susceptible.

**Table 1.**
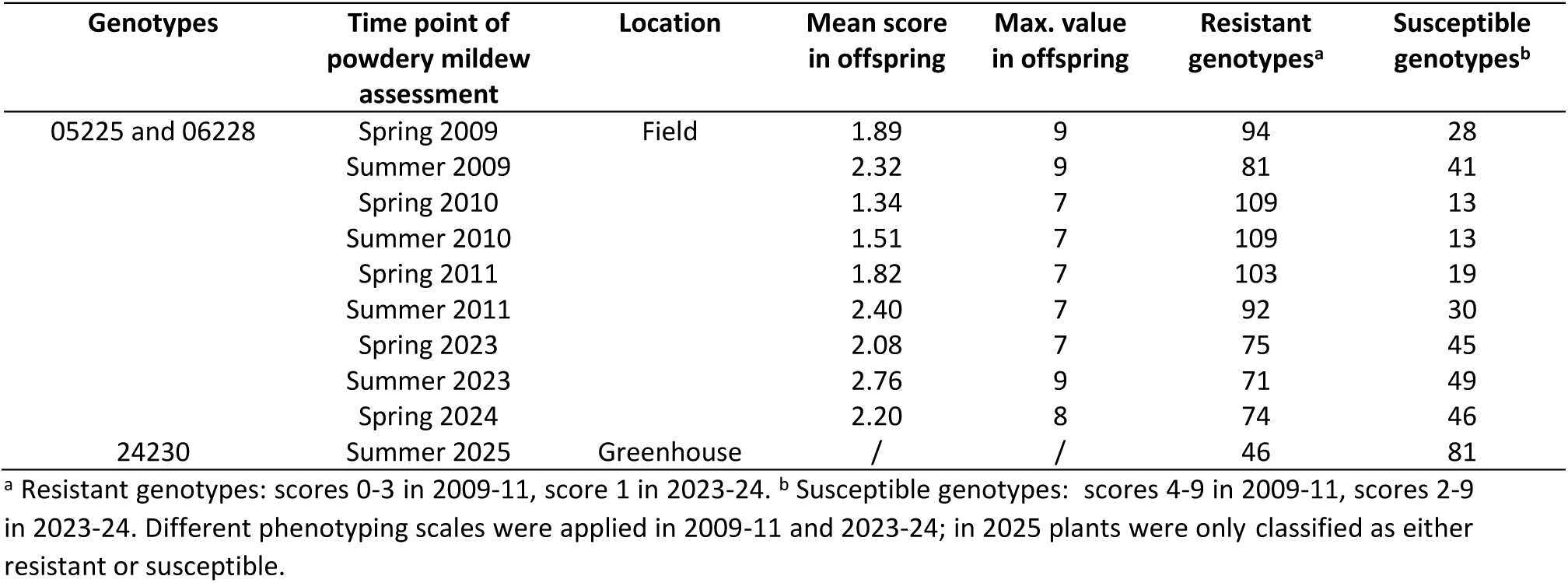
Overview of phenotyping results from the ‘Idared’ × *Malus baccata* ‘Jackii’ F_1_ populations in the field and the greenhouse.

### Genetic linkage map of *Mb*j haplotype 1

A genetic linkage map was constructed for *Mb*j haplotype 1 (HT1), comprising 17 linkage groups with a total length of 1,071 centimorgans (cM). From the 324,420 non-imputed single-nucleotide polymorphism (SNP) markers, 261,000 were excluded as they were either homozygous in *Mb*j or were inconsistent across the four *Mb*j replicates. From the remaining 63,420 SNPs (19.6%), those with > 10% missing data in the progeny and chi-square values ≥ 10 were discarded, resulting in 29,245 SNP markers. From the imputed data of these markers, a total of 2,043 with the largest inter-marker distances were loaded into JoinMap 5, and after excluding identical markers, this resulted in 948 SNP markers being distributed across the 17 LGs. Of the 63 simple sequence repeat (SSR) markers selected from the HiDRAS website (HiDRAS 2025), 58 could be assigned to the 17 LGs, with three of them displaying multilocus alleles. Of the seven newly developed SSRs, five were mapped to LG 10, and one each to LG 5 and LG 7. Table S1 provides an overview of the number of markers per LG and their genetic lengths. Figure S1 shows the 17 LGs, including marker names and their respective positions in cM.

### QTL analysis reveals the major *Plbj* resistance locus and a novel minor locus

Kruskal-Wallis (KW) analysis using the phenotypic data of the primary F_1_ mapping population, together with the constructed genetic linkage map, revealed a consistent and significant association of markers on LG 10 and LG 5 with resistance to powdery mildew across all the phenotyped time points in 2009, 2010, 2011, 2023, and 2024. The highest KW values for markers on LG 10 ranged from 51.15 to 109.04 (df = 1 or 3; significance level: *p* < 0.0001), whereas those for LG 5 ranged from 9.67 to 17.96 (df = 1; significance level: *p* < 0.005). Interval mapping confirmed significant associations on both LGs 10 and 5. Table 2 presents the markers with the highest LOD scores on these LGs. The major QTL on LG 10 showed LOD scores ranging from 16.34 to 35.73, while the minor QTL on LG 5 had scores between 2.49 and 4.63. A permutation test showed that the QTL on LG 10 consistently exceeded the genome-wide threshold, whereas the QTL on LG 5 surpassed the genome-wide significance threshold only with data from spring 2009. In spring 2011 and 2023, as well as in the summers of 2009-11, LOD scores on LG 5 only exceeded the chromosome-wide significance threshold. Figure 1 shows the LOD score profiles for the identified QTLs on LGs 10 and 5. Whereas the QTL on LG 10 explained up to 74% of the phenotypic variance, the QTL on LG 5 explained between 9.1% and 16.0% of the phenotypic variance. Newly developed SSR markers LKSSRchr10_1478 and LKSSRchr10_2718 were found to flank the QTL region on LG 10, with LKSSRchr10_1998 and LKSSRchr10_2318B being highly linked to the QTL peak. LKSSRchr5_Mbj2 was associated with resistance conferred by the minor QTL on LG 5.

**Figure 1.**
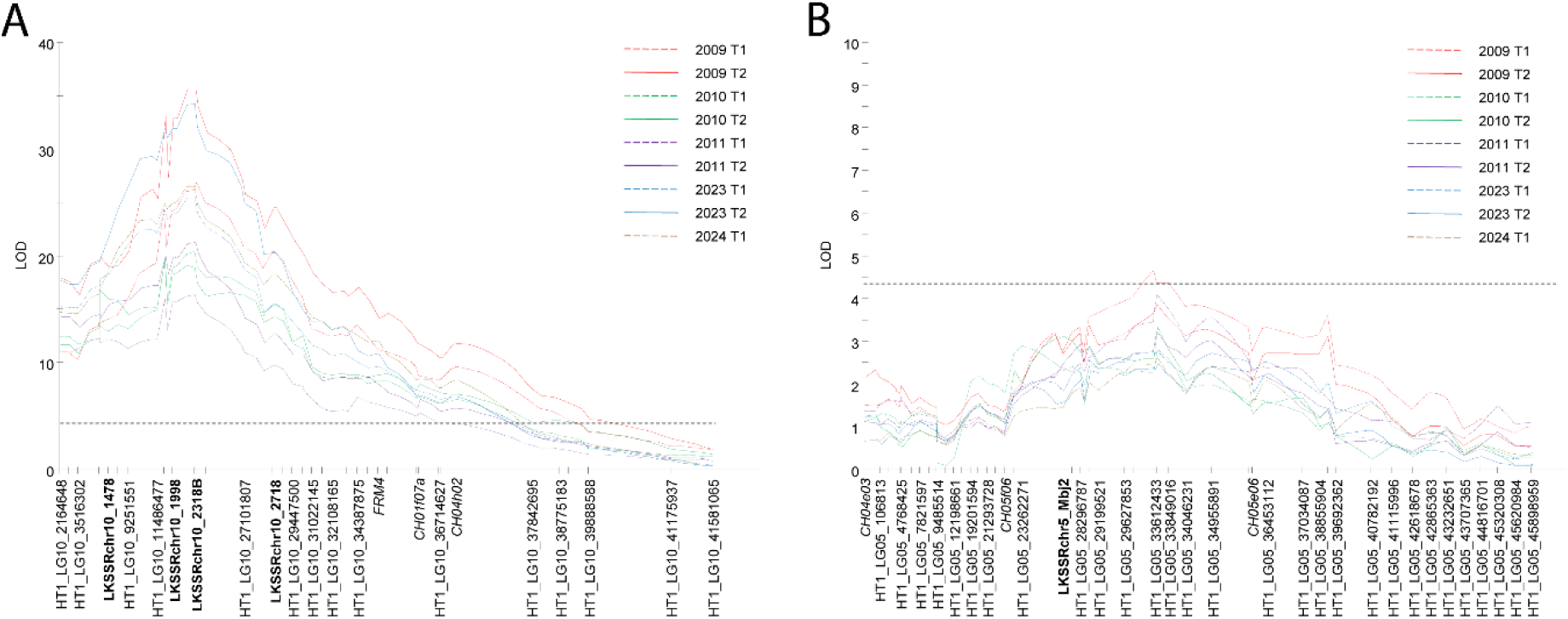
LOD score profiles of the identified QTLs on linkage group 10 (A) and 5 (B) of *Malus baccata* ‘Jackii’, derived from powdery mildew infection data at various time points, indicating the approximate position of *Plbj.* T1: time point 1 (spring), T2: time point 2 (summer). The dotted line represents the genome-wide significance threshold. SSRs developed in this project are shown in bold and SSRs selected from the HiDRAS website (HiDRAS 2025) in italics. For a better readability, only a subset of markers is presented.

**Table 2.** Markers with the highest LOD scores on linkage groups 10 and 5.

| Time point of powdery mildew assessment | Linkage group | Marker with highest LOD score | LOD score | Explained variance (%) | Average phenotypic score with/without resistance allele | Significance with permutation test at a 95% confidence level |
| --- | --- | --- | --- | --- | --- | --- |
| Spring 2009 | 10 | HT1_LG10_25523367 | 26.92 | 63.8 | 0.39 / 3.92 | Genome-wide |
| Summer 2009 |  | LKSSRchr10_2318B | 35.73 | 74.0 | 0.43 / 4.96 | Genome-wide |
| Spring 2010 |  | LKSSRchr10_2318B | 20.39 | 53.7 | 0.19 / 2.88 | Genome-wide |
| Summer 2010 |  | LKSSRchr10_1978 | 19.87 | 52.8 | 0.43 / 2.84 | Genome-wide |
| Spring 2011 |  | LKSSRchr10_2318B | 16.34 | 46.0 | 0.78 / 3.37 | Genome-wide |
| Summer 2011 |  | LKSSRchr10_2318B | 21.24 | 55.1 | 1.22 / 4.22 | Genome-wide |
| Spring 2023 |  | LKSSRchr10_2318B | 25.50 | 62.4 | 1.03 / 3.56 | Genome-wide |
| Summer 2023 |  | LKSSRchr10_2318B | 34.27 | 73.2 | 1.00 / 5.28 | Genome-wide |
| Spring 2024 |  | LKSSRchr10_2318B | 26.56 | 63.9 | 1.05 / 3.90 | Genome-wide |
| Spring 2009 | 5 | HT1_LG05_33612433 | 4.63 | 16.0 | 2.50 / 4.48<sup>a</sup> | Genome-wide |
| Summer 2009 |  | HT1_LG05_31890599 | 3.85 | 13.5 | 3.93 / 5.42<sup>a</sup> | Chromosome-wide only |
| Spring 2010 |  | HT1_LG05_23262271 | 2.92 | 10.5 | 2.13 / 3.27<sup>a</sup> | Not significant |
| Summer 2010 |  | HT1_LG05_31890599 | 4.00 | 14.0 | 2.18 / 3.34<sup>a</sup> | Chromosome-wide only |
| Spring 2011 |  | HT1_LG05_31890599 | 4.09 | 14.3 | 2.43 / 3.65<sup>a</sup> | Chromosome-wide only |
| Summer 2011 |  | HT1_LG05_31890599 | 3.24 | 11.5 | 3.62 / 4.25<sup>a</sup> | Chromosome-wide only |
| Spring 2023 |  | HT1_LG05_31890599 | 2.78 | 10.1 | 2.87 / 3.93<sup>a</sup> | Chromosome-wide only |
| Summer 2023 |  | HT1_LG05_31890599 | 2.49 | 9.1 | 4.56 / 5.66<sup>a</sup> | Not significant |
| Spring 2024 |  | HT1_LG05_31890599 | 2.62 | 9.6 | 3.12 / 4.24<sup>a</sup> | Not significant |

<sup>a</sup> For linkage group 5 only offspring without *Plbj* were considered to avoid distortion in the average phenotypic score.

### *Plbj* is located between markers LKSSRchr10_1998 and LKSSRchr10_2318B

In the primary mapping population in the field (122 F_1_ individuals), all plants lacking the resistance-associated SSR allele of LKSSRchr10_2318B were phenotyped as susceptible to powdery mildew in the 2023-24 dataset, whereas only one individual carrying the resistance alleles of all five newly developed SSRs on LG 10 was also phenotyped as susceptible. Phenotyping in 2009-11 identified seven individuals carrying susceptibility alleles of these five SSRs that were nonetheless phenotyped as resistant. In the secondary population (127 greenhouse-grown F_1_ individuals), three individuals also showed genotype-phenotype incongruence as they carried only resistance-associated SSR alleles but were phenotyped as susceptible, whereas all individuals lacking resistance-associated marker alleles were phenotyped as susceptible. Among the 45 *Mb*j seedlings (pollinated by F_1_ individuals derived from ‘Idared’ × *Mb*j), five individuals with recombinations between LKSSRchr10_1478 and LKSSRchr10_2718 were identified. One showed visible powdery mildew symptoms, while the remaining four were presumed resistant; however, confirmation under higher powdery mildew pressure is required. Mapping of *Plbj* as a single qualitative trait using the 2023-24 phenotypic dataset of the primary mapping population positioned the locus between SSR markers LKSSRchr10_1998 and LKSSRchr10_2318B (Figure 2). To validate this, 26 individuals with recombinations between LKSSRchr10_1478 and LKSSRchr10_2718 from different populations were examined. A comparison of these recombinants (Figure 3) revealed that genotypes possessing the resistance alleles of LKSSRchr10_1998 and LKSSRchr10_2318B exhibited a resistant phenotype, whereas genotypes lacking the resistance alleles at both markers exhibited a susceptible phenotype. In contrast, the presence of only one resistance allele at either of the two markers could result in either a resistant or a susceptible phenotype due to recombination events. Individuals 06228-068, 24230-043 and Jackii-OA-115 indicate that *Plbj* lies downstream of LKSSRchr10_1998, whereas 24230-039 and Jackii-OA-81 suggest it is upstream of LKSSRchr10_2318B. Taken together, these findings delimit *Plbj* to the interval between LKSSRchr10_1998 and LKSSRchr10_2318B.

**Figure 2.**
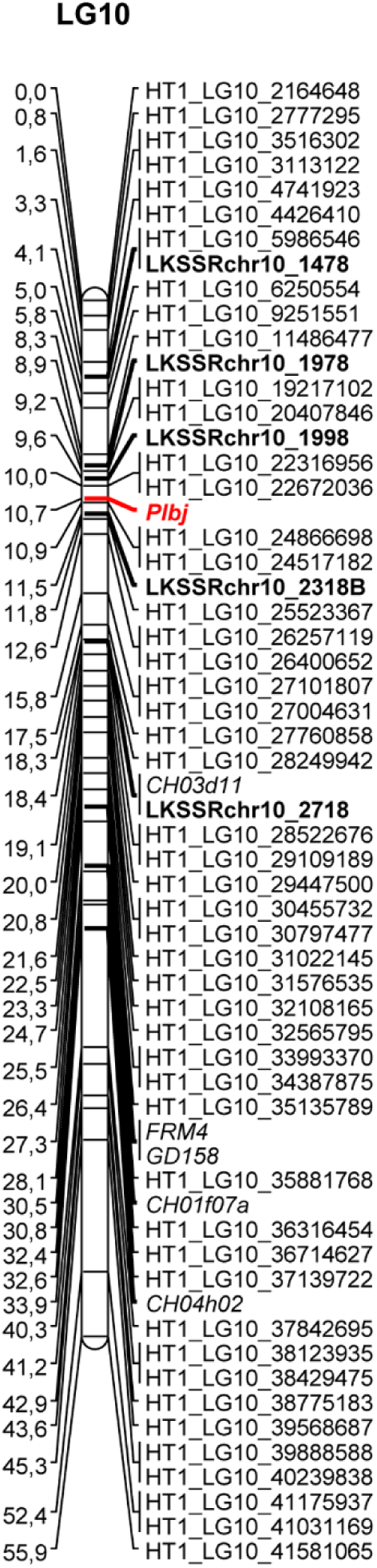
Genetic linkage map of linkage group 10 of haplotype 1 of *Malus baccata* ‘Jackii’, showing the approximate position of *Plbj*. SSRs developed in this project are shown in bold, SSRs selected from the HiDRAS website (HiDRAS 2025) in italics, and *Plbj* is indicated with a bold red italic label.

**Figure 3.**
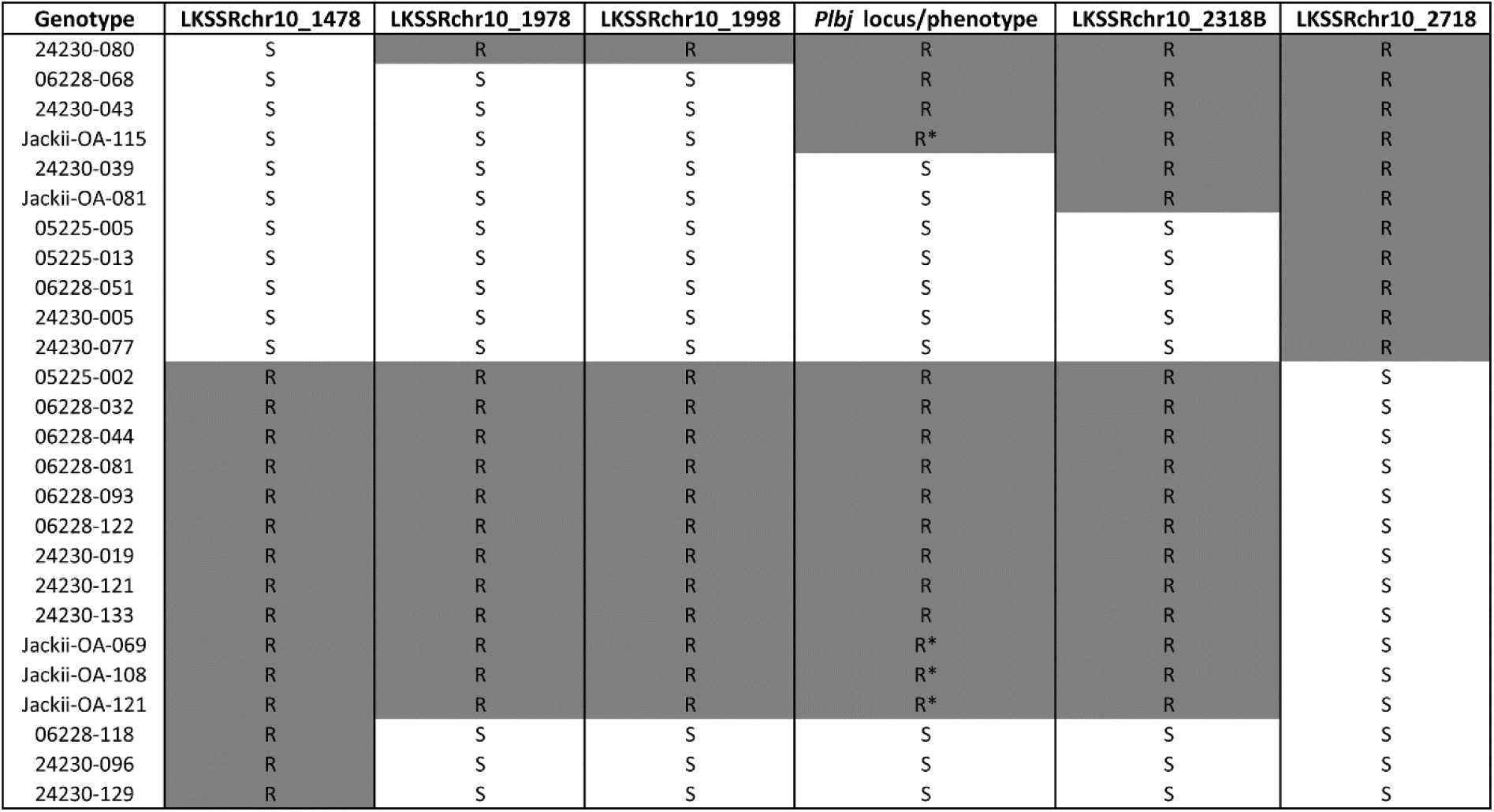
Graphical representation of the *Plbj* region based on SSR markers. Twenty-six individuals show recombination events between SSR markers LKSSRchr10_1478 and LKSSRchr10_2718, suggesting that *Plbj* is located between the SSR markers LKSSRchr10_1998 and LKSSRchr10_2318B. Genotypes 05225 and 06228 are recombinant individuals from the primary F_1_ mapping population maintained in the field, whereas genotypes 24230 (secondary F_1_ population) and Jackii-OA (*Malus baccata* ‘Jackii’ seedlings pollinated by F_1_ individuals derived from ‘Idared’ × *Malus baccata* ‘Jackii’) are maintained in the greenhouse. R: resistance allele, S: susceptibility allele. For the *Plbj* locus/phenotype R denotes a resistant and S denotes a susceptible phenotype; an asterisk indicates resistance presumed due to low powdery mildew pressure.

### SNP and BLAST-supported SSR marker analysis identifies resistance-associated haplotypes

A comparison between the expected PCR product sizes of the newly developed SSRs and those observed in the fragment length analysis revealed differences of up to five bp. Nevertheless, the size differences between resistance- and susceptibility-associated alleles within each marker were consistent with the expected values. When phenotypic data were included, it became evident that *Plbj* is linked to the marker alleles assigned to HT1. In contrast, lower average disease scores in individuals lacking *Plbj* were associated with allele 111 from LKSSRchr5_Mbj2, indicating that the minor QTL on LG 5 is located on HT2. SSR LKSSRchr10_1438, designed for LG 10 based on the GDDH13 genome (Daccord et al. 2017), amplified fragments that were not linked to powdery mildew resistance and instead mapped to LG 7 in *Mb*j. An overview of the results for the six newly developed SSRs linked to powdery mildew resistance is provided in Table S2. The assignment of *Plbj* and the minor QTL on LG 5 to their respective haplotypes was further confirmed by examining selected SNP markers linked to the two resistance loci. Based on the known parental alleles and the segregation pattern in the F₁ progeny, the resistance-associated SNPs contributed by *Mb*j could be assigned to the corresponding haplotypes. As shown in Table 3, *Plbj* is located on HT1, while the minor QTL on LG 5 is located on HT2.

**Table 3.** SNP marker-based identification of resistance-associated haplotypes in *Malus baccata* ‘Jackii’.

| Marker | SNP-alleles in 'Idared' | SNP-allele in *Malus baccata* 'Jackii' haplotype 1 | SNP-allele in *Malus baccata* 'Jackii' haplotype 2 | Resistance-associated SNP-allele combination in offspring | Resistance-associated SNP-allele in *Malus baccata* 'Jackii' |
| --- | --- | --- | --- | --- | --- |
| HT1_LG10_22316956 | GG | C | G | CG | C |
| HT1_LG10_24866698 | CC | T | C | TC | T |
| HT1_LG10_25523367 | TT | T | C | TT | T |
| HT1_LG05_29627853 | GG | A | G | GG<sup>a</sup> | G |
| HT1_LG05_33612433 | AA | A | T | AT<sup>a</sup> | T |
| HT1_LG05_33849016 | AA | G | A | AA<sup>a</sup> | A |

<sup>a</sup> For linkage group 5 only offspring without *Plbj* were considered to avoid distortion in identifying the resistance-associated SNP-allele combination.

### Validation of KASP markers through comparison to SSR markers

The newly developed Kompetitive Allele Specific PCR (KASP) markers KASP_HT1_LG10_20042456 and KASP_HT1_LG10_23962775, located between LKSSRchr10_1478 and LKSSRchr10_1978, and between LKSSRchr10_1998 and LKSSRchr10_2318B, respectively, showed complete concordance with the SSR marker data. No apparent double recombination events were observed between the flanking SSR markers in any individual of the primary F_1_ mapping population. Specifically, for KASP_HT1_LG10_20042456, the resistance-associated allele was consistently linked to the T_Hex_ allele, and for KASP_HT1_LG10_23962775, resistance was associated with the G_Hex_ allele. Moreover, the KASP-assays were tested on nine founders of apple ‘Idared’, ‘Golden Delicious’, ‘Granny Smith’, ‘Delicious’, ‘Cox Orange’, ‘Jonathan’, ‘Mcintosh’, ‘Braeburn’ and ‘Gala’ and all of them consistently exhibited the FAM-labeled C-allele linked to susceptibility to powdery mildew in *Mb*j.

### Resistance gene candidates within the *Plbj* locus on HT1

The region of interest, defined as the interval between markers LKSSRchr10_1998 and LKSSRchr10_2318B on HT1 (specifically from position 21,391,236 to 24,678,521 bp, spanning 3,287,286 bp), contains a total of 209 annotated genes (Table S3). Among these, 29 genes were identified as the most likely resistance gene candidates (Table S4).

## Discussion

Powdery mildew is one of the most important fungal diseases in apple, and its impact may further increase in the future (García-Gómez et al. 2024; Strickland et al. 2021). At the same time, demand for reduced pesticide use is steadily growing, making genetic host resistance increasingly important in sustainable apple production. Pyramiding different resistance genes not only has the potential to increase durability but can even reduce disease severity beyond that of the single most effective resistance gene (Mundt 2018). Consequently, substantial efforts in apple breeding aim to combine high fruit quality with multiple resistance genes targeting either the same pathogen or different diseases simultaneously (Baumgartner et al. 2015; Kellerhals et al. 2013). For powdery mildew, genotypes carrying *Pl-1* and *Pl-2* have been bred (Kellerhals et al. 2013). However, both of these resistance genes have already been overcome when deployed individually (Caffier and Laurens 2005; Caffier and Parisi 2007; Kellerhals et al. 2013; Krieghoff 1995). This highlights the urgent need to incorporate additional, more robust sources of powdery mildew resistance into apple cultivars.

In the present study, a genetic linkage map of *Mb*j HT1 was generated, spanning a total length of 1,071 cM, which is comparable to previously published genetic linkage maps in apple (Celton et al. 2009; Emeriewen et al. 2020; Norelli et al. 2017). Most of the SNP markers were ordered consistently with their physical positions, supporting the overall accuracy of the map. Moreover, nearly all mapped SSR markers corresponded well with positions reported on the HiDRAS website (HiDRAS 2025). When comparing the physical positions of SNP markers of each chromosome to the position of the corresponding linkage group, it became evident that the SNPs typically started within the first few megabases and extended toward the chromosome ends, while gaps remain in some intermediate regions. One notable exception was LG 7, where the first mapped SNP marker was located at 31 Mb, and the linkage group itself measured only 19.1 cM. This suggests that the proximal region of chromosome 7 is a recombination-poor region in the studied population, likely resulting in the exclusion of markers up to 31 Mb during linkage map construction. Importantly, this linkage map allowed the detection of loci associated with powdery mildew resistance on LG 5 and LG 10.

Because powdery mildew in apple is caused by various pathogen strains (Gañán-Betancur et al. 2021; Lesemann et al. 2004; Urbanietz and Dunemann 2005), phenotypic evaluation under natural, high-disease-pressure field conditions over multiple years can be considered a robust method to assess the durability and effectiveness of a resistance gene. Therefore, the result of this study, namely that *Plbj* was consistently detected in an F_1_ population from 2009-11 and 2023-24 under field conditions without fungicide application, assessed by two different persons with different assessment scales, exhibits the durability and heritability of this resistance. LOD scores on LG 10 exceeded the genome-wide significance threshold at every phenotyping time point. The percentage of explained variance varied between the phenotyping time points, likely due to differences in disease pressure and weather conditions, but reached up to 74%. Since similarly high LOD scores (> 25) and explained variances (> 65%) have been reported for major resistance genes against fire blight (Broggini et al. 2014; Emeriewen et al. 2014; Emeriewen et al. 2018; Fahrentrapp et al. 2013; Peil et al. 2007), it can be hypothesised that *Plbj* also represents a major monogenic resistance gene. Therefore, the precise identification and delimitation of the genomic region harbouring *Plbj* is crucial. In this study, the candidate region could be narrowed down to a 3,287,286 bp interval through genetic mapping. Since this region has been sequenced (Pfeifer et al. 2025, preprint), an initial prediction of 29 potential resistance genes was possible. This represents an important first step toward the functional analysis of *Plbj*. Nevertheless, future fine-mapping approaches using more recombinants, combined with the development of additional markers in the candidate region, are expected to further improve resolution and narrow down the locus. Moreover, the recent availability of the *Podosphaera leucotricha* genome will facilitate future research on the fungus itself as well as host-pathogen interactions (Gañán et al. 2020), especially once specific resistance genes in *Mb*j have been identified.

The strong effect of *Plbj* was clearly detectable in all years of the study. However, a few genotype-phenotype incongruences were observed. One genotype that carried the resistance marker alleles of the five newly develepod SSRs on LG 10 was nevertheless scored as susceptible, with scores of three and four at two field phenotyping time points in 2023 and 2024. As this genotype was scored as resistant in most years, a misclassification in those two years, possibly due to confusion with susceptible neighbouring trees, appears plausible. Conversely, three greenhouse-grown individuals from the 24230 population carried the resistance marker alleles of *Plbj,* but still showed noticeable powdery mildew symptoms in 2025. Possible explanations include mutations in the resistance gene itself, the presence of modifier genes suppressing resistance expression, or stress-induced susceptibility caused by limited plant spacing, reduced sunlight and partial spider mite infestation in the greenhouse. Furthermore, greenhouse conditions are markedly different from field conditions and have an influence on powdery mildew susceptibility (Jeger et al. 1986). Interestingly, it has already been described that genotypes with *Pl-m* appearing susceptible under artificial conditions still exhibit resistance in the field (Bus et al. 2010), suggesting that similar effects may explain these observations.

In addition to *Plbj*, a minor QTL on LG 5 HT2 was identified, showing chromosome-wide significance in three out of five scoring years. This QTL has not been previously reported by Dunemann and Schuster (2009). It remains unclear whether this locus represents a single gene or polygenic resistance. However, as the phenotypic variance explained by this locus was only up to 16%, compared to 74% for *Plbj*, and the LOD peak was noticeably flatter, it can be hypothesised that resistance at this locus is polygenic. Polygenic resistance is often more durable than monogenic major resistance genes (Parlevliet 2002). Therefore, minor QTLs such as the one detected on LG 5 should not be underestimated. Indeed, it has been reported that QTLs for powdery mildew resistance on LGs 1, 8, 10, 14, and 17 are sometimes identified only in specific years, with phenotypic variation explained ranging from 5.1 to 19.5%, in contrast to stable QTLs on LGs 2 and 13, for which the phenotypic variation explained ranges from 7.5 to 27.4% across years (Calenge and Durel 2006). In our study, a QTL peak on LG 5 was observed in all years, even when it did not consistently exceed the chromosome-wide significance threshold in interval mapping. This, together with a significant KW association, suggests that the underlying resistance effect was present across all years but varied in strength between years.

Molecular markers are nowadays crucial for MAS, allowing selection based solely on genotypic information. For various powdery mildew resistance genes in apple, molecular markers have been reported (Bus et al. 2010; Dunemann et al. 2007; Dunemann and Schuster 2009; García-Gómez et al. 2024; Gardiner et al. 2003; James and Evans 2004; Luo et al. 2019; Markussen et al. 1995; Seglias and Gessler 1997). However, some of these resistance genes, such as *Pl-1* (Kellerhals et al. 2013; Krieghoff 1995) and *Pl-2* (Caffier and Laurens 2005; Caffier and Parisi 2007), have already been overcome and for others, like *Plbj,* the previously most closely linked marker was an AFLP/SCAR marker. In contrast, SSR and KASP markers are now more commonly used. In this study, we employed the published haplotype-resolved genome sequence of *Mb*j (Pfeifer et al. 2025, preprint), which facilitated the development of five SSR and two KASP markers linked to the resistance locus on LG 10 and one SSR linked to the resistance on LG 5. KASP markers are becoming increasingly important, as their analysis is relatively cheap and rapid, requiring only a real-time PCR machine rather than a costly capillary electrophoresis genetic analyser. The markers reported here can now be readily used in apple breeding programmes to efficiently select progenies with *Plbj* resistance. The potential presence of both a major monogenic resistance locus (*Plbj*) and a minor polygenic QTL in *Mb*j makes this genotype highly attractive for breeding, as it may allow the combination of both types of resistance, each with its respective advantages, within a single donor genotype. Moreover, *Mb*j has previously been shown to exhibit resistance to apple scab (Gygax et al. 2004), several strains of fire blight (Vogt et al. 2013; Wöhner et al. 2018), and tolerance to *Diplocarpon coronariae* (Wöhner et al. 2021). Taken together, *Mb*j represents an exceptionally valuable donor for future apple resistance breeding, which is gaining in importance for sustainable apple production.

## Materials and methods

### Plant material

The apple cultivar ‘Idared’ and the apple genotype *Mb*j are susceptible and resistant to powdery mildew, respectively (Figure 4). A cross between ‘Idared’ and *Mb*j resulted in 122 F_1_ individuals (05225 and 06228 genotypes), which served as the primary mapping population for this study. These progenies are cultivated in the experimental field of the Julius Kühn-Institut (JKI) in Dresden-Pillnitz, Germany, without fungicide protection. Additionally, our study included a recently developed secondary population consisting of 127 individuals from the same cross combination (24230 genotypes), as well as 45 seedlings derived from *Mb*j pollinated by F_1_ individuals of ‘Idared’ × *Mb*j (designated as Jackii-OA), all of which are maintained under greenhouse conditions without fungicide application.

**Figure 4.**
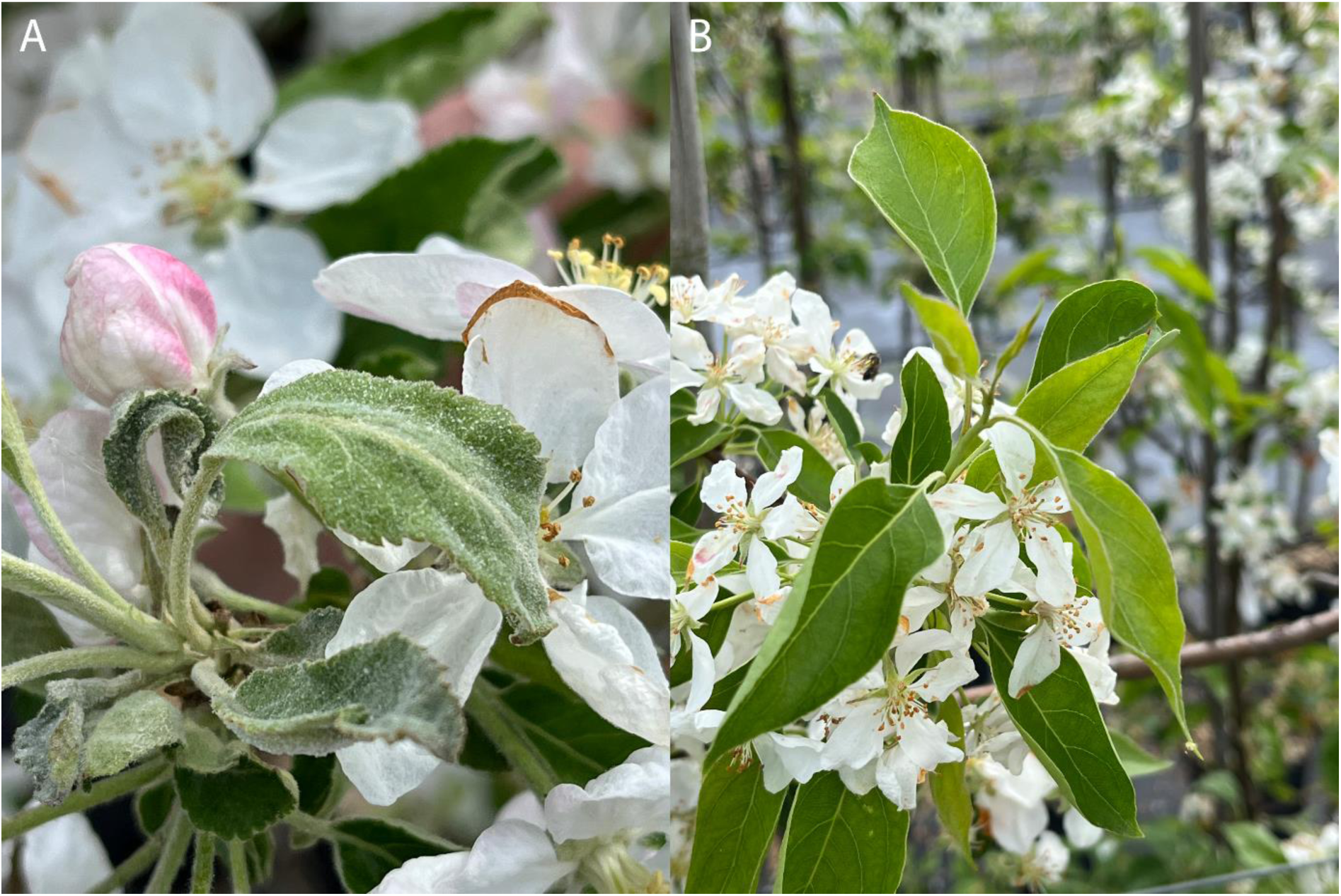
Symptoms of powdery mildew on ‘Idared’ (A), and absence of symptoms on *Malus baccata* ‘Jackii’ (B).

### Powdery mildew phenotyping

Phenotyping of natural powdery mildew infestation was conducted for the primary mapping population (05225 and 06228 genotypes) in the field during the spring and summer of 2009, 2010, 2011 and 2023. In 2024, assessments were performed only in spring, as the trees were pruned afterwards and the prevalence of powdery mildew post-pruning was too low for a reliable evaluation. Table 4 shows the phenotyping scales for the assessment of powdery mildew infestation. The scale applied in 2023 and 2024 was published by Lateur et al. (2022). For 2009-11, plants with powdery mildew infestation scores of zero to three were classified as resistant, whereas those with scores of four to nine were classified as susceptible. For 2023-24, plants with a score of one were considered resistant, while those with scores from two to nine were considered susceptible. Plants grown in the greenhouse (24230 and Jackii-OA genotypes) were classified only as either resistant or susceptible, without any intermediate categories, based on the presence or absence of disease symptoms resulting from natural infection.

**Table 4.** Used phenotyping scales for powdery mildew infection.

| Score | Phenotyping scale 2009-11 | Phenotyping scale 2023-24 (Lateur et al. 2022) |
| --- | --- | --- |
| 0 | No visible symptom | not applicable |
| 1 | Very few sporulating dots | No visible symptom (0%) |
| 2 | Very few to few sporulating dots | One or very few organs affected, detectable on close scrutiny of the tree (0-1%) |
| 3 | Up to 25% of the tree affected by infected leaves/shoots | Infected organs readily apparent but without important consequences for the tree (1-5%) |
| 4 | Intermediate rating | Intermediate rating |
| 5 | Up to 50% of the tree affected by infected leaves/shoots | Primary mildew widespread over the branches, inducing the infection of a substantial part of the crown ($\pm 25\%$) |
| 6 | Intermediate rating | Intermediate rating |
| 7 | Up to 75% of the tree affected by infected leaves/shoots | Heavy infection; half of the organs are badly affected ($\pm 50\%$) |
| 8 | Intermediate rating | Intermediate rating |
| 9 | Up to 100% of the tree affected by infected leaves/shoots | Crown completely affected, nearly all top of the organs are infected ($> 90\%$) |

### tGBS genotyping and SNP identification

Young leaves of the 122 F_1_ individuals of the primary mapping population along with four replicates of both parents were harvested, lyophilised, and sent to Data2Bio (Ames, IA, USA) for DNA extraction and tunable genotyping-by-sequencing (tGBS) analysis (Ott et al. 2017) using the restriction enzyme *Bsp*1286I and an Illumina HiSeq X instrument (Illumina, Inc., San Diego, CA, USA), according to the company’s specifications. tGBS genotyping and SNP identification were performed as described by Pfeifer et al. (2025, preprint). Briefly, quality-trimmed sequence reads, excluding regions with a PHRED score ≤ 15, were aligned to HT1 of the *Mb*j genome using GSNAP (Wu and Nacu 2010). Only confidently mapped reads that aligned to a unique location in HT1 were used for SNP identification. For homozygous SNPs, the most common allele had to be supported by at least 80% of all aligned reads at a given position and confirmed by a minimum of five unique reads. For heterozygous SNPs, the two most common alleles each had to be supported by at least 30% of the aligned reads at that position and confirmed by at least five unique reads. The minimum calling rate for SNPs across the population was set to ≥ 50%, and the minor allele frequency had to be ≥ 10%. Finally, SNPs lacking a sufficient number of reads to make genotype calls were imputed using Beagle v5.4 (Browning et al. 2018). Similar empirical parameters, which aim to minimise false positive and false negative SNP calls, have already been applied in wild *Malus* species (Emeriewen et al. 2020) and other plants (Li et al. 2018; Zheng et al. 2018).

### SSR marker sourcing, development and genotyping

Since tGBS-derived SNPs are not easily transferable to other sample sets, we sourced SSR markers from the literature to serve as anchor markers for the construction of genetic linkage maps (Celton et al. 2009; Emeriewen et al. 2014; Emeriewen et al. 2017; Hemmat et al. 2003; Hokanson et al. 1998; Liebhard et al. 2002; Silfverberg-Dilworth et al. 2006; Vinatzer et al. 2004; Yamamoto et al. 2002a; Yamamoto et al. 2002b). Initially, 115 SSRs were selected to be distributed across all 17 chromosomes of *Malus*, from the HiDRAS website (HiDRAS 2025) and tested for polymorphism in the parents ‘Idared’ and *Mb*j and a subset of six offspring. Of the 71 SSRs showing polymorphisms in *Mb*j, a total of 63 were used in this study (Table S5). In addition, seven new SSRs were developed to flank the resistance loci. Therefore, SSR motifs were searched for in the genome sequences of GDDH13 v1.1 (Daccord et al. 2017) and *Mb*j (Pfeifer et al. 2025, preprint), in the regions identified by preliminary QTL mapping and primer pairs (Table S6) were designed using Primer3web v4.1.0 (Koressaar and Remm 2007; Untergasser et al. 2007). The 122 F_1_ individuals (05225 and 06228 genotypes) grown in the field and both parents were genotyped with the 63 SSRs selected from the HiDRAS website (Table S5) and the seven newly developed SSR markers (Table S6). The 127 F_1_ individuals (24230 genotypes) cultivated in the greenhouse, as well as the 45 *Mb*j seedlings (Jackii-OA genotypes), were genotyped with only five of the newly developed SSR markers.

### DNA isolation, PCR and fragment analysis

DNA was extracted from leaves using the DNeasy Plant Mini Kit (Qiagen, Hilden, Germany) according to the manufacturer’s protocol. DNA quantification was performed with the NanoDrop One^C^ spectrophotometer (Thermo Fisher Scientific Inc., Waltham, MA, USA). Multiplex-PCR was conducted using the Type-it Microsatellite PCR Kit (Qiagen, Hilden, Germany). The PCR reaction mix consisted of 1 μl primer or primer mix (each primer at a concentration of 1 pmol/µl), 1 μl ddH_2_O, 5 μl Type-it Multiplex PCR Master Mix, 1 μl Q-solution, and 2 μl DNA (10 ng/μl). The PCR conditions were as follows: initial denaturation at 95°C for 5 minutes, followed by 32 cycles of 95°C for 1 minute, 60°C for 1 minute and 30 seconds, and 72°C for 1 minute, with a final elongation step at 60°C for 30 minutes. The PCR products were then diluted 1:100 with ddH_2_O, and 1μl of this dilution was mixed with 9 μl of ABI-solution (a mixture of 1 ml Hi-Di formamide and 6 μl GeneScan-600 LIZ size standard, both from Applied Biosystems, Waltham, MA, USA). The samples were then denatured at 95°C for 5 minutes prior to analysis using the Applied Biosystems 3500xL Genetic Analyzer (Applied Biosystems, Waltham, MA, USA). GeneMapper Software v6 (Applied Biosystems, Waltham, MA, USA) was used to visualise and analyse the SSR alleles.

### Construction of genetic linkage maps

Prior to genetic linkage map construction, the 324,420 non-imputed SNP markers for the *Mb*j HT1 were filtered. In the first step, all SNP markers that did not yield identical results across all four replicates of ‘Idared’ and *Mb*j, were excluded. SNP markers with more than 10% missing values in the progeny were also excluded. For hk×hk and nn×np markers, chi-square values were calculated, and only markers with values below 10 were retained. For each linkage group, SNPs were ranked by physical inter-marker distance and all markers with inter-marker distances of 100 kb or greater were included and at least 120 markers per group were selected. From the selected SNPs, the imputed genotypic data along with the 70 SSRs were used for the construction of the genetic map of *Mb*j using JoinMap 5 (Van Ooijen 2018). The final map of *Mb*j was calculated after the exclusion of identical markers and correcting implausible double-recombinations using the regression mapping algorithm and Kosambi’s mapping function. Linkage maps were visualised using MapChart (Voorrips 2002).

### QTL analysis

The SNP and SSR genotypic data of the primary F_1_ mapping population (05225 and 06228 genotypes), the final genetic map of *Mb*j together with multi-year phenotypic datasets, were used to determine genotype-phenotype associations and conduct QTL analysis using MapQTL 5 (Van Ooijen 2004). Kruskal-Wallis analysis was applied to identify markers significantly associated with the phenotypic data, whereas interval mapping was used to localise the corresponding QTL intervals. A permutation test at a 95% confidence level was conducted to assess the significance of identified QTLs by determining the LOD threshold at both the genome-wide and chromosome-wide levels (Van Ooijen 2004).

### Mapping resistance as a single qualitative trait

To map resistance as a single qualitative trait, named *Plbj*, phenotypic data from 2023-24 for the 122 F_1_ individuals in the field (05225 and 06228 genotypes) were transformed for each individual into resistant (score one) or susceptible (scores two to nine). These data were then added to the other molecular markers on LG 10, and the genetic map was calculated using JoinMap 5 (Van Ooijen 2018).

### Assigning resistance-linked markers to haplotypes of *Mb*j

To identify the resistance-associated haplotypes of *Plbj* and the minor QTL on LG 5, two independent approaches were applied. In the first approach, the primer sequences of the newly developed SSR markers were aligned to the genome sequences of both haplotypes of *Mb*j (Pfeifer et al. 2025, preprint) using the Basic Local Alignment Search Tool (BLAST; Altschul et al. 1990) and CLC Main Workbench 25.0 (Qiagen, Venlo, Netherlands). Expected PCR product sizes in base pairs were calculated by considering the distance between the outermost primer positions in the haplotypes and accounting for any additional bases present in the primers but absent from the assembled genome. The expected PCR product sizes were then compared with the fragment sizes observed in the fragment length analysis. In the second approach, the alleles of nn×np SNP markers from both parents and the primary mapping population (122 F_1_ individuals) were examined. Based on the inheritance patterns and the assumption that *Mb*j is the resistance donor, the SNP alleles associated with resistance were identified. In both approaches, resistance-associated markers were determined by comparing the mean disease scores of the 122 F_1_ individuals for the respective allele combinations and assigning resistance to the markers associated with lower average disease severity. For the assignment of the minor QTL on LG 5, only individuals lacking *Plbj* were taken into account to avoid distortion of the disease score average caused by *Plbj* when identifying the resistance-associated markers.

### KASP marker development and genotyping

Based on the genetic linkage map, two KASP markers were developed in proximity to *Plbj*. The SNP positions of these KASP markers are reflected in their names: KASP_HT1_LG10_20042456 and KASP_HT1_LG10_23962775. Primer sequences for these KASP markers are listed in Table S7. KASP genotyping was performed using a reaction mix consisting of the KASP assay mix (a mixture of two allele-specific forward primers and one common reverse primer), KASP-TF V4.0 2× Master Mix (LGC Group, Teddington, England) and DNA. Reactions were run on a CFX96 Touch Real-Time PCR Detection System (Bio-Rad Laboratories, Inc., Hercules, CA, USA). The 10 µl KASP reaction contained 5 μl 2× KASP master mix, 0.14 μl KASP assay mix, 1 μl DNA (10 ng/μl) and 3.86 μl ddH_2_O. PCR was performed under the following conditions: an initial activation at 94°C for 15 minutes, followed by 10 cycles of denaturation at 94°C for 20 seconds and annealing/elongation for 1 minute with a temperature gradient from 61 to 55°C (decreasing 0.6°C per cycle), and then 26 cycles of 94°C for 20 seconds and 55°C for 1 minute and finally, a 1-minute step at 37°C for the read stage. Data analysis was performed with CFX Manager v3.1 (Bio-Rad Laboratories, Inc., Hercules, CA, USA). DNA from the cultivars ‘Idared’, ‘Golden Delicious’, ‘Granny Smith’, ‘Delicious’, ‘Cox Orange’, ‘Jonathan’, ‘Mcintosh’, ‘Braeburn’, ‘Gala’ and *Mb*j were used as control for the validation of the KASP markers.

### Identification of resistance gene candidates in the genome of *Mb*j

Resistance geme candidates were identified based on annotation data published by Pfeifer et al. (2025, preprint). Genes that were annotated with an NB-ARC domain (PF00931), leucine-rich repeat N-terminal domain (PF08263), TIR domain (PF01582), Rx N-terminal domain (PF18052) or associated with the Gene Ontology terms defence response (GO:0006952), response to other organism (GO:0051707) or protein kinase activity (GO:0004672), as well as proteins whose names contained the term disease resistance, were considered the most likely resistance gene candidates.

## Acknowledgements

Parts of this work were supported by the Federal Ministry of Agriculture, Food and Regional Identity by decision of the Parliament of the Federal Republic of Germany via the Federal Office for Agriculture and Food (BLE) under the innovation support programmes 281D108X21 and 281D109A21. Language editing was supported using DeepL Write and ChatGPT (OpenAI), which were used for language improvement only.

## Author Contributions

Conception: Ofere Francis Emeriewen, Andreas Peil, Thomas Wöhner and Henryk Flachowsky. Strategy and design: Matthias Pfeifer, Ofere Francis Emeriewen, Andreas Peil and Thomas Wöhner. Analyses and writing: Matthias Pfeifer. Plant material: Matthias Pfeifer and Andreas Peil. Phenotyping: Matthias Pfeifer and Andreas Peil. Primer design: Ofere Francis Emeriewen and Leonard Kurzweg. Marker analyses: Matthias Pfeifer, Ofere Francis Emeriewen, Leonard Kurzweg, Buist Muçaj and Tom Burkhardt. Genetic map construction: Matthias Pfeifer, Andreas Peil, Ofere Francis Emeriewen and Leonard Kurzweg. QTL analyses: Matthias Pfeifer, Ofere Francis Emeriewen, Andreas Peil and Leonard Kurzweg. Genomic analyses: Matthias Pfeifer and Thomas Wöhner. Funding: Andreas Peil and Thomas Wöhner. Supervision: Ofere Francis Emeriewen, Andreas Peil, Thomas Wöhner and Henryk Flachowsky. Revision: All authors. All authors read and approved the final manuscript.

## Data Availability Statement

The data underlying this article will be shared on reasonable request to the corresponding author.

## Conflicts of Interests

The authors declare no conflicts of interests.

## Supplementary information

**Figure S1** Genetic linkage maps of haplotype 1 of *Malus baccata* ‘Jackii’. SSRs developed in this project are shown in bold, SSRs selected from the HiDRAS website (HiDRAS 2025) in italics, and *Plbj* is indicated with a bold red italic label (see separate supplementary file).

**Table S1.** Number of markers and genetic length (cM) of the 17 linkage groups of *Malus baccata* ‘Jackii’ haplotype 1.

| Linkage group | SNP marker | SSR marker | Length (cM) |
| --- | --- | --- | --- |
| 1 | 47 | 5 | 63.6 |
| 2 | 70 | 4 | 67.1 |
| 3 | 58 | 5 | 78.6 |
| 4 | 58 | 2 | 50.8 |
| 5 | 62 | 4 | 66.6 |
| 6 | 53 | 2 | 65.9 |
| 7 | 24 | 2 | 19.1 |
| 8 | 55 | 4 | 56.3 |
| 9 | 51 | 4 | 60.7 |
| 10 | 50 | 10 | 55.9 |
| 11 | 53 | 3 | 69.6 |
| 12 | 69 | 1 | 64.3 |
| 13 | 51 | 7 | 68.1 |
| 14 | 61 | 2 | 59.9 |
| 15 | 65 | 3 | 107.4 |
| 16 | 67 | 5 | 63.6 |
| 17 | 54 | 6 | 53.5 |
| Total | 948 | 69 | 1071.0 |

**Table S2.** Summary of newly developed SSR markers linked to powdery mildew resistance.

| Marker | Linkage group | Predicted genomic position in haplotype 1 / 2 | Expected PCR product size in bp of haplotype 1 / 2<sup>a</sup> | Observed size in fragment length analysis (bp)<sup>a</sup> |
| --- | --- | --- | --- | --- |
| LKSSRchr5_Mbj2 | 5 | 25,400,896-25,400,994 / 25,183,631-25,183,741 | 99 / 111 | 99 / 111 |
| LKSSRchr10_1478 | 10 | 17,120,061-17,120,242 / 16,946,103-16,946,288 | **182** / 186 | **177** / 181 |
| LKSSRchr10_1978 | 10 | 21,155,277-21,155,449 / 21,035,950-21,036,126 | **175** / 179 | **176** / 180 |
| LKSSRchr10_1998 | 10 | 21,391,017-21,391,236 / 21,254,415-21,254,632 | **220** / 218 | 215 / **217** |
| LKSSRchr10_2318B | 10 | 24,678,521 - 24,678,663 / 24,412,181-24,412,327 | **143** / 147 | **143** / 147 |
| LKSSRchr10_2718 | 10 | 28,170,148-28,170,275 / 27,895,197-27,895,319 | **128** / 123 | 119 / **124** |

<sup>a</sup> Resistance-associated alleles are shown in bold.

**Table S3** Annotated genes in the region of interest in haplotype 1 of linkage group 10 in *Malus baccata* ‘Jackii’ (see separate supplementary file).

**Table S4** Identified resistance gene candidates for *Plbj* (see separate supplementary file).

**Table S5.** Previously published SSR markers selected from the HiDRAS website (HiDRAS 2025)

| Reference | Marker names |
| --- | --- |
| Celton et al. 2009 | NZmsCN943067, NZmsCO754252, NZmsEB137525, NZmsMDAJ1681 |
| Emeriewen et al. 2014 | FRM4 |
| Emeriewen et al. 2017 | FRMb251 |
| Hemmat et al. 2003 | GD153, GD158 |
| Hokanson et al. 1998 | GD96, GD142, GD147 |
| Liebhard et al. 2002 | CH01e01, CH01f03b, CH01f07a, CH01f09, CH01h02, CH01h10, CH02a03, CH02b03b, CH02b10, CH02c06, CH02d08, CH02d12, CH02f06, CH02g01, CH02g09, CH02h11a, CH03a08, CH03b10, CH03d07, CH03d11, CH03e03, CH03g07, CH03g12, CH04e03, CH04f10, CH04h02, CH05b06, CH05c07, CH05e06, CH05f06, CH05g08 |
| Silfverberg-Dilworth et al. 2006 | AU223657-SSR, Hi01d05, Hi02c06, Hi02c07, Hi03a10, Hi03d06, Hi03e04, Hi03g06, Hi04b12, Hi04d02, Hi04e04, Hi04f09, Hi04g05, Hi05b09, Hi07b02, Hi07h02, Hi08f12, MDAJ761-SSR |
| Vinatzer et al. 2004 | CH-Vf1 |
| Yamamoto et al. 2002a | KA4b |
| Yamamoto et al. 2002b | NH033b |

**Table S6.**
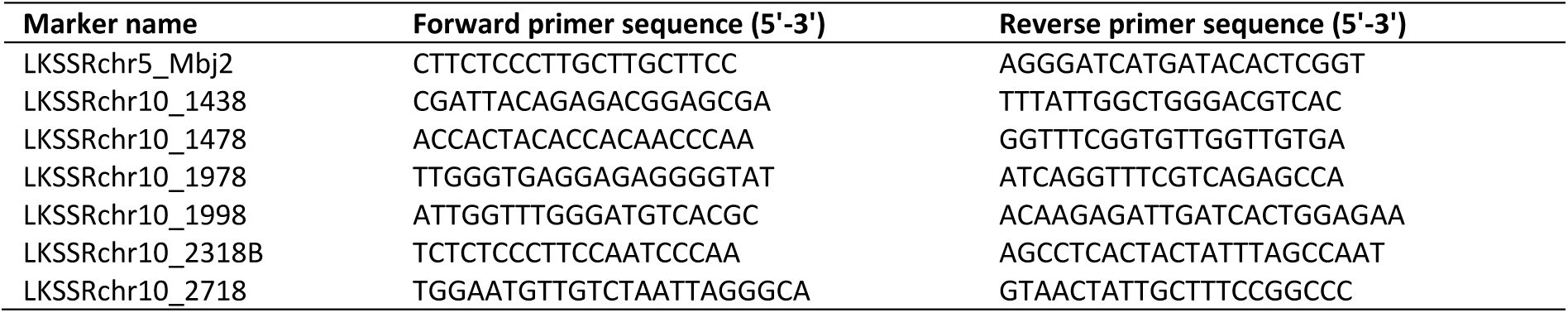
Newly developed SSR markers with primer sequences.

**Table S7.**
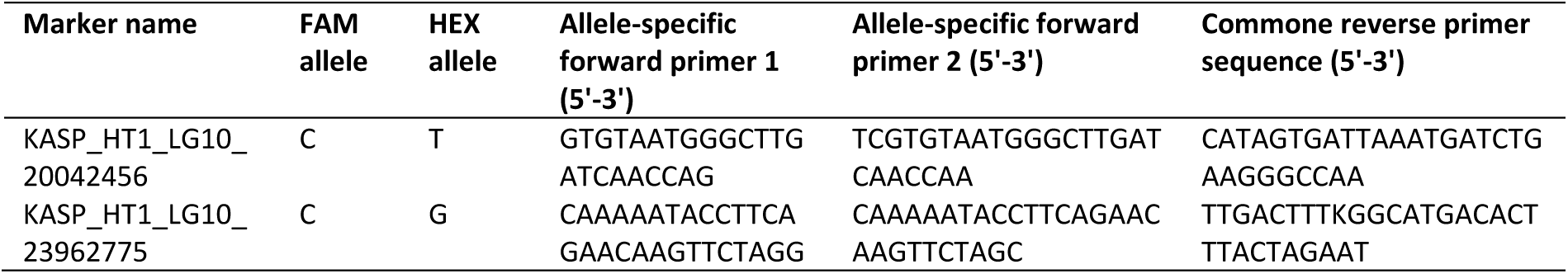
Overview of developed KASP markers.

## Notes

### Competing Interest Statement

The authors have declared no competing interest.

## References

1. Altschul, S. F., W. Gish, W. Miller, E. W. Myers, and D. J. Lipman. 1990. “Basic local alignment search tool.“ Journal of Molecular Biology 215: 403–410.

2. Baumgartner, I. O., A. Patocchi, J. E. Frey, A. Peil, and M. Kellerhals. 2015. “Breeding Elite Lines of Apple Carrying Pyramided Homozygous Resistance Genes Against Apple Scab and Resistance Against Powdery Mildew and Fire Blight.“ Plant Molecular Biology Reporter 33: 1573–1583.

3. Broggini, G. A. L., T. Wöhner, et al. 2014. “Engineering fire blight resistance into the apple cultivar ‘Gala’ using the *FB_MR5* CC-NBS-LRR resistance gene of *Malus* × *robusta* 5.“ Plant Biotechnology Journal 12: 728–733.

4. Browning, B. L., Y. Zhou, and S. R. Browning. 2018. “A One-Penny Imputed Genome from Next-Generation Reference Panels.“ The American Journal of Human Genetics 103: 338–348.

5. Bus, V. G. M., H. C. M. Bassett, et al. 2010. “Genome mapping of an apple scab, a powdery mildew and a woolly apple aphid resistance gene from open-pollinated Mildew Immune Selection.“ Tree Genetics & Genomes 6: 477–487.

6. Caffier, V., and F. Laurens. 2005. “Breakdown of *Pl2*, a major gene of resistance to apple powdery mildew, in a French experimental orchard.“ Plant Pathology 54: 116–124.

7. Caffier, V., and L. Parisi. 2007. “Development of apple powdery mildew on sources of resistance to *Podosphaera leucotricha*, exposed to an inoculum virulent against the major resistance gene *Pl-2*.“ Plant Breeding 126: 319–322.

8. Calenge, F., and C.-E. Durel. 2006. “Both stable and unstable QTLs for resistance to powdery mildew are detected in apple after four years of field assessments.“ Molecular Breeding 17: 329–339.

9. Celton, J.-M., D. S. Tustin, D. Chagné, and S. E. Gardiner. 2009. “Construction of a dense genetic linkage map for apple rootstocks using SSRs developed from *Malus* ESTs and *Pyrus* genomic sequences.“ Tree Genetics & Genomes 5: 93–107.

10. Coyier, D. L. 1974. “Heterothallism in the Apple Powdery Mildew Fungus, *Podosphaera leucotricha*.“ Phytopathology 64: 246–248.

11. Daccord, N., J.-M. Celton, et al. 2017. “High-quality de novo assembly of the apple genome and methylome dynamics of early fruit development.“ Nature genetics 49: 1099–1106.

12. Dayton, D. F. 1977. “Genetic Immunity to Apple Mildew Incited by *Podosphaera leucotricha*.“ American Society for Horticultural Science 12: 225–226.

13. Dunemann, F., A. Peil, A. Urbanietz, and T. Garcia-Libreros. 2007. “Mapping of the apple powdery mildew resistance gene *Pl1* and its genetic association with an NBS-LRR candidate resistance gene.“ Plant Breeding 126: 476–481.

14. Dunemann, F., and M. Schuster. 2009. “Genetic characterization and mapping of the major powdery mildew resistance gene *Plbj* from *Malus baccata jackii*.“ Acta Horticulturae 814: 791–798.

15. Emeriewen, O., K. Richter, et al. 2014. “Identification of a major quantitative trait locus for resistance to fire blight in the wild apple species *Malus fusca*.“ Molecular Breeding 34: 407–419.

16. Emeriewen, O. F., A. Peil, K. Richter, E. Zini, M.-V. Hanke, and M. Malnoy. 2017. “Fire blight resistance of *Malus* ×*arnoldiana* is controlled by a quantitative trait locus located at the distal end of linkage group 12.“ European Journal of Plant Pathology 148: 1011–1018.

17. Emeriewen, O. F., K. Richter, et al. 2020. “Construction of a dense genetic map of the *Malus fusca* fire blight resistant accession MAL0045 using tunable genotyping-by-sequencing SNPs and microsatellites.“ Scientific Reports 10: 16358.

18. Emeriewen, O. F., K. Richter, et al. 2018. “Towards map-based cloning of *FB_Mfu10*: identification of a receptor-like kinase candidate gene underlying the *Malus fusca* fire blight resistance locus on linkage group 10.“ Molecular Breeding 38: 106.

19. Fahrentrapp, J., G. A. L. Broggini, et al. 2013. “A candidate gene for fire blight resistance in *Malus* × *robusta* 5 is coding for a CC–NBS–LRR.“ Tree Genetics & Genomes 9: 237–251. DOI: 10.1007/s11295-012-0550-3.

20. Gallott, J. C., R. C. Lamb, and H. S. Aldwinckle. 1985. “Resistance to Powdery Mildew from Some Small-fruited *Malus* Cultivars.“ American Society for Horticultural Science 20: 1085–1087.

21. Gañán, L., R. A. White III, M. L. Friesen, T. L. Peever, and A. Amiri. 2020. “A Genome Resource for the Apple Powdery Mildew Pathogen *Podosphaera leucotricha*.“ Phytopathology 110: 1756–1758.

22. Gañán-Betancur, L., T. L. Peever, K. Evans, and A. Amiri. 2021. “High Genetic Diversity in Predominantly Clonal Populations of the Powdery Mildew Fungus *Podosphaera leucotricha* from U.S. Apple Orchards.“ Applied and Environmental Microbiology 87: e00469–21.

23. García-Gómez, B. E., S. Bühlmann-Schütz, M. Hodel, A. Patocchi, and M. J. Aranzana. 2024. “Development of SNP markers for the powdery mildew resistance gene *Pl1* in apple.“ Acta Horticulturae 1412: 299–306.

24. Gardiner, S. E., J. Murdoch, et al. 2003. “Candidate resistance genes from an EST database prove a rich source of markers for major genes conferring resistance to important apple pests and diseases.“ Acta Horticulturae 622: 141–151.

25. Garibaldi, A., G. Gilardi, and M. L. Gullino. 2005. “First Report of Powdery Mildew Caused by *Podosphaera leucotricha* on *Photinia* × *fraserii* in Italy.“ Plant Disease 89: 1362.

26. Gygax, M., L. Gianfranceschi, R. Liebhard, M. Kellerhals, C. Gessler, and A. Patocchi. 2004. “Molecular markers linked to the apple scab resistance gene *Vbj* derived from *Malus baccata jackii*“. Theoretical and Applied Genetics 109: 1702–1709.

27. Hanke, M.-V., H. Flachowsky, A. Peil, and O. F. Emeriewen. 2020. “Chapter 19.3—Malus × domestica Apple.“ In Biotechnology of Fruit and Nut Crops, edited by R. E. Litz, F. Pliego-Alfaro, and J. I. Hormaza, 440–473. UK: CAB International.

28. Hemmat, M., N. F. Weeden, and S. K. Brown. 2003. “Mapping and Evaluation of *Malus* ×*domestica* Microsatellites in Apple and Pear.“ jashs 128: 515–520.

29. HiDRAS: High-quality Disease Resistant Apples for a Sustainable Agriculture (2025). Available at: https://sites.unimi.it/camelot/hidras/index.php. [Accessed 14 March 2025].

30. Hokanson, S. C., A. K. Szewc-McFadden, W. F. Lamboy, and J. R. McFerson. 1998. “Microsatellite (SSR) markers reveal genetic identities, genetic diversity and relationships in a Malus×domestica borkh. core subset collection.“ Theoretical and Applied Genetics 97: 671–683.

31. James, C. M., and K. M. Evans. 2004. “ Identification of molecular markers linked to the mildew resistance genes *Pl-d* and *Pl-w* in apple.“ Acta Horticulturae 663: 123–128.

32. Jeger, M. J., D. J. Butt, and A. A. J. Swait. 1986. “Components of resistance of apple to powdery mildew (*Podosphaera leucotricha*).“ Plant Pathology 35: 477–490.

33. Kellerhals, M., I. O. Baumgartner, L. Leumann, J. E. Frey, and A. Patocchi. 2013. “Progress in Pyramiding Disease Resistances in Apple Breeding.“ Acta Horticulturae 976: 487–491.

34. Knight, R. L. and F. H. Alston. 1968. “Sources of field immunity to mildew (*Podosphaera leucotricha*) in apple.“ Canadian Journal of Genetics and Cytology 10: 294–298.

35. Koressaar, T., and M. Remm. 2007. “Enhancements and modifications of primer design program Primer3.“ Bioinformatics 23: 1289–1291.

36. Krieghoff, O. 1995. “Entwicklung einer In-vitro-Selektionsmethode auf Resistenz von Malus- Genotypen gegenüber Podosphaera leucotricha (Ell. et Ev.) Salm. und In-vitro-Differenzierung von Virulenzunterschieden des Erregers.“ Doctoral Dissertation, Humboldt-Universität zu Berlin, Germany.

37. Kusch, S., J. Qian, A. Loos, F. Kümmel, P. D. Spanu, and R. Panstruga. 2024. “Long-term and rapid evolution in powdery mildew fungi.“ Molecular Ecology 33: e16909.

38. Lateur, M., E. Dapen, et al. 2022. “ECPGR Characterization and Evaluation Descriptors for Apple Genetic Resources.“ European Cooperative Programme for Plant Genetic Resources, Rome, Italy.

39. Lesemann, S., A. Urbanietz, and F. Dunemann. 2004. “ Determining population variation of apple powdery mildew at the molecular level.“ Acta Horticulturae 663: 199–204.

40. Lesemann, S. S., S. Schimpke, F. Dunemann, and H. B. Deising. 2006. “Mitochondrial heteroplasmy for the cytochrome b gene Controls the level of strobilurin resistance in the apple powdery mildew fungus *Podosphaera leucotricha* (Ell. & Ev.) E.S. Salmon. “Journal of Plant Diseases and Protection 113: 259–266.

41. Li, T., J. Qu, et al. 2018. “Genetic characterization of inbred lines from Shaan A and B groups for identifying loci associated with maize grain yield.“ BMC Genetics 19: 63.

42. Liebhard, R., L. Gianfranceschi, et al. 2002. “Development and characterisation of 140 new microsatellites in apple (*Malus* x *domestica* Borkh.).“ Molecular Breeding 10: 217–241.

43. Luo, F., P. Sandefur, K. Evans, and C. Peace. 2019. “A DNA test for routinely predicting mildew resistance in descendants of crabapple ‘White Angel’.“ Molecular Breeding 39: 33.

44. Markussen, T., J. Krüger, H. Schmidt, and F. Dunemann. 1995. “Identification of PCR-based markers linked to the powdery-mildew-resistance gene *Pl*1 from *Malus robusta* in cultivated apple*.“* Plant Breeding 114: 530–534.

45. Minnis, A. M., A. Y. Rossman, D. L. Clement, M. K. Malinoski, and K. K. Rane. 2010. “First Report of Powdery Mildew Caused by *Podosphaera leucotricha* on Callery Pear in North America.“ Plant Disease 94: 279.

46. Mundt, C. C. 2018. “Pyramiding for Resistance Durability: Theory and Practice.“ Phytopathology 108: 792–802.

47. Mwanza, E. J. M., S. K. Waithaka, and S. A. Simons. 2001. “First Report of Powdery Mildew Caused by *Podosphaera leucotricha* on *Prunus africana* in Kenya.“ Plant Disease 85: 1285.

48. Norelli, J. L., M. Wisniewski, et al. 2017. “Genotyping-by-sequencing markers facilitate the identification of quantitative trait loci controlling resistance to *Penicillium expansum* in *Malus sieversii*.“ PlOS One 12: e0172949.

49. Ott, A., S. Liu, et al. 2017. “tGBS® genotyping-by-sequencing enables reliable genotyping of heterozygous loci.“ Nucleic Acids Research 45: e178.

50. Parlevliet, J. E. 2002. “Durability of resistance against fungal, bacterial and viral pathogens; present situation.“ Euphytica 124: 147–156.

51. Peil, A., T. Garcia-Libreros, et al. 2007. “Strong evidence for a fire blight resistance gene of *Malus robusta* located on linkage group 3.“ Plant Breeding 126: 470–475.

52. Pessina, S., D. Angeli, et al. 2016. “The knock-down of the expression of *MdMLO19* reduces susceptibility to powdery mildew (*Podosphaera leucotricha*) in apple (*Malus domestica*).“ Plant Biotechnology Journal 14: 2033–2044.

53. Pessina, S., L. Palmieri, et al. 2017. “Frequency of a natural truncated allele of *MdMLO19* in the germplasm of *Malus domestica*.“ Molecular Breeding 37: 7.

54. Pfeifer, M., O. F. Emeriewen, et al. 2025. “High-quality haplotype-resolved genome assembly and annotation of Malus baccata ‘Jackii’.“ Preprint available at: https://www.biorxiv.org/content/10.1101/2025.07.27.667097v1. [Accessed 1 August 2025].

55. Schuster, M. 2000. “ Genetics of powdery mildew resistance in *Malus* species.“ Acta Horticolturae 538: 593–595.

56. Seglias, N. P., and C. Gessler. 1997. “Genetics of apple powdery mildew resistance from *Malus zumi* (*Pl2*).“ IOBC-WPRS Bulletins 20: 195–208.

57. Silfverberg-Dilworth, E., C. L. Matasci, et al. 2006. “Microsatellite markers spanning the apple (*Malus* x *domestica* Borkh.) genome.“ Tree Genetics & Genomes 2: 202–224.

58. Stankiewicz-Kosyl, M., E. Pitera, and S. W. Gawronski. 2005. “Mapping QTL involved in powdery mildew resistance of the apple clone U 211.“ Plant Breeding 124: 63–66.

59. Strickland, D. A., K. T. Hodge, and K. D. Cox. 2021. “An Examination of Apple Powdery Mildew and the Biology of *Podosphaera leucotricha* from Past to Present.“ Plant Health Progress 22: 421–432.

60. Strickland, D. A., J. P. Spychalla, E. van Zoeren, M. R. Basedow, D. J. Donahue, and K. D. Cox. 2023. “Assessment of Fungicide Resistance via Molecular Assay in Populations of Podosphaera leucotricha, Causal Agent of Apple Powdery Mildew, in New York.“ Plant Disease 107: 2606–2612.

61. Takamatsu, S. 2013. “Molecular phylogeny reveals phenotypic evolution of powdery mildews (Erysiphales, Ascomycota).“ Journal of General Plant Pathology 79: 218–226.

62. Untergasser, A., I. Cutcutache, et al. 2012. “Primer3—new capabilities and interfaces.“ Nucleic Acids Research 40: e115.

63. Urbanietz, A., and F. Dunemann. 2005. “Isolation, identification and molecular characterization of physiological races of apple powdery mildew (*Podosphaera leucotricha*).“ Plant Pathology 54: 125– 133.

64. Van Ooijen, J. W. 2004. “MapQTL 5, Software for the mapping of quantitative trait loci in experimental populations.“ Kyazma B. V., Wageningen, Netherlands.

65. Van Ooijen, J. W. 2018. “JoinMap5, Software for the calculation of genetic linkage maps in experimental populations of diploid species.“ Kyazma B.V., Wageningen, Netherlands.

66. Vielba-Fernández, A., Á. Polonio, L. Ruiz-Jiménez, A. de Vicente, A. Pérez-García, and D. Fernández-Ortuño. 2020. “Fungicide Resistance in Powdery Mildew Fungi.“ Microorganisms 8: 1431.

67. Vinatzer, B. A., A. Patocchi, S. Tartarini, L. Gianfranceschi, S. Sansavini, and C. Gessler. 2004. “Isolation of two microsatellite markers from BAC clones of the *Vf* scab resistance region and molecular characterization of scab-resistant accessions in *Malus* germplasm*.“ Plant Breeding 123: 321–326.

68. Visser, T., and J. J. Verhaegh. 1980. “Resistance to powdery mildew (*Podosphaera leucotricha*) of apple seedlings growing under glasshouse and nursery conditions.“ *Proceedings of the Eucarpia meeting of tree fruit breeding*, Angers, 1979, 111–120.

69. Vogt, I., T. Wöhner, et al. 2013. “Gene-for-gene relationship in the host-pathogen system *Malus* × *robusta* 5-*Erwinia amylovora*.“ New Phytologist 197: 1262–1275.

70. Voorrips, R. E. 2002. “MapChart: Software for the Graphical Presentation of Linkage Maps and QTLs.“ Journal of Heredity 93: 77–78.

71. Wöhner, T. W., K. Richter, et al. 2018. “Inoculation of *Malus* genotypes with a set of *Erwinia amylovora* strains indicates a gene-for-gene relationship between the effector gene *eop1* and both *Malus floribunda* 821 and *Malus* ‘Evereste’.“ Plant Pathology 67: 938–947.

72. Wöhner, T., O. F. Emeriewen, and M. Höfer. 2021. “Evidence of apple blotch resistance in wild apple germplasm (*Malus* spp.) accessions.“ European Journal of Plant Pathology 159: 441–448.

73. Wu, T. D., and S. Nacu. 2010. “Fast and SNP-tolerant detection of complex variants and splicing in short reads.“ Bioinformatics 26: 873–881.

74. Yamamoto, T., T. Kimura, et al. 2002a. “Simple sequence repeats for genetic analysis in pear.“ Euphytica 124: 129–137.

75. Yamamoto, T., T. Kimura, M. Shoda, Y. Ban, T. Hayashi, and N. Matsuta. 2002b. “Development of microsatellite markers in the Japanese pear (*Pyrus pyrifolia* Nakai).“ Molecular Ecology Notes 2: 14– 16.

76. Yoder, K. S. 2000. “Effect of Powdery Mildew on Apple Yield and Economic Benefits of Its Management in Virginia.“ Plant Disease 84: 1171–1176.

77. Zheng, Z., Z. Sun, et al. 2018. “Genetic Diversity, Population Structure, and Botanical Variety of 320 Global Peanut Accessions Revealed Through Tunable Genotyping-by-Sequencing.“ Scientific Reports 8: 14500.

