## Supplementary figures and images for "Genetic mapping, marker development, and identification of candidate genes for powdery mildew resistance in *Malus baccata* ‘Jackii’"

### Figure S1

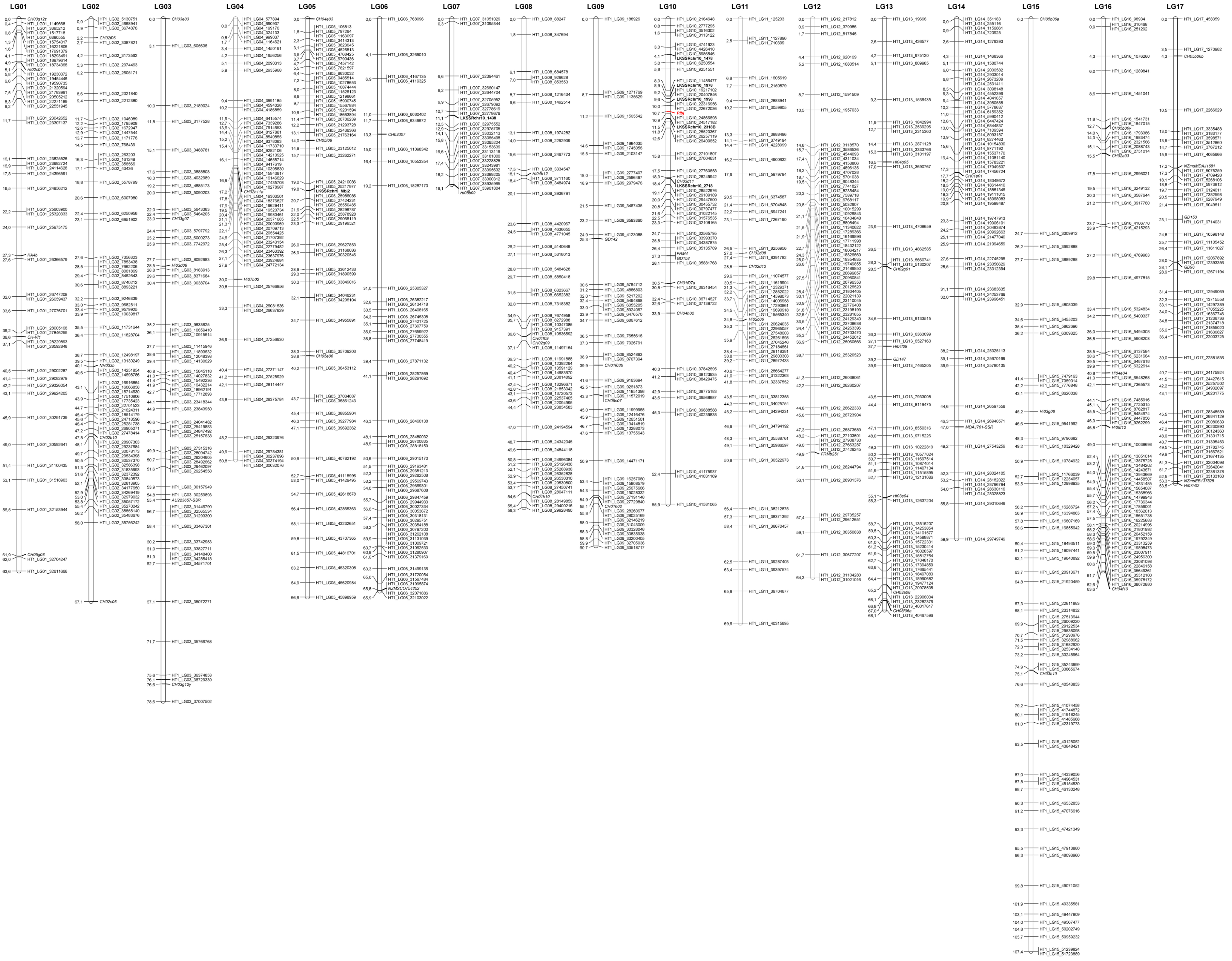
